# Characterization of the motor cortex transcriptome supports microgial-related key events in amyotrophic lateral sclerosis

**DOI:** 10.1101/2020.02.07.938662

**Authors:** Oriol Dols-Icardo, Victor Montal, Sònia Sirisi, Gema López-Pernas, Laura Cervera-Carles, Marta Querol-Vilaseca, Laia Muñoz, Olivia Belbin, Daniel Alcolea, Laura Molina-Porcel, Jordi Pegueroles, Janina Turón-Sans, Rafael Blesa, Alberto Lleó, Juan Fortea, Ricard Rojas-García, Jordi Clarimón

## Abstract

Amyotrophic lateral sclerosis (ALS) is a devastating neurodegenerative disease characterized by the degeneration of upper and lower motor neurons. A major neuropathological finding in ALS is the coexistence of glial activation and aggregation of the phosphorylated transactive response DNA-binding protein 43-kDa (pTDP43) in the motor cortex at the earliest stages of the disease. Despite this, the transcriptional alterations associated with these pathological changes in this major vulnerable brain region have yet to be fully characterized. Here, we have performed massive RNA sequencing of the motor cortex of ALS (n=11) and healthy controls (HC; n=8). We report extensive RNA expression alterations at gene and isoform levels, characterized by the enrichment of neuroinflammatory and synapse related pathways. The assembly of gene co-expression modules confirmed the involvement of these two principal transcriptomic changes, and showed a strong negative correlation between them. Furthermore, cell-type deconvolution using human single-nucleus RNA sequencing data as reference demonstrated that microglial cells are overrepresented in ALS compared to HC. Importantly, we also show for the first time in the human ALS motor cortex, that microgliosis is mostly driven by the increased proportion of a microglial subpopulation characterized by gene markers overlapping with the recently described disease associated microglia (DAM). Using immunohistochemistry, we further evidenced that this microglial subpopulation is overrepresented in ALS and that variability in pTDP43 aggregation among patients negatively correlates with the proportion of microglial cells. In conclusion, we report that neuroinflammatory changes in ALS motor cortex are dominated by microglia which is concomitant with a reduced expression of postsynaptic transcripts, in which DAM might have a prominent role. Microgliosis therefore represents a promising avenue for therapeutic intervention in ALS.

## Introduction

Amyotrophic lateral sclerosis (ALS) is a neurodegenerative disease characterized by the degeneration of upper and lower motor neurons leading to progressive muscle weakness, wasting and paralysis that result in death within a few years from disease onset. The aberrant cytoplasmic aggregation and phosphorylation of the 43-kDa transactive response DNA-binding protein (pTDP-43) represents the main pathological hallmark in the majority of ALS cases at postmortem evaluation and is known to occur in a sequential manner beginning in the motor cortex (1). Although the causes that lead to non-genetic forms of ALS (sporadic ALS) have yet to be fully determined, the presence of neuroinflammation is a consistent feature in the central nervous system of affected subjects (2). Microglia are resident macrophages that belong to the immune system of the central nervous system (CNS), where, as a result of disease conditions, change their gene expression profile to adapt and acquire a reactive state (3). Recently, a study using single-cell RNA sequencing of brain tissue from an ALS mouse model reported that a specific microglial subpopulation, known as disease associated microglia (DAM), was the cell-type responsible for ALS-related microgliosis and that is enhanced by triggering receptor expressed on myeloid cells 2 (TREM2) (4). In support of this scenario, some chitinase proteins, which are known markers of microglial activation (5, 6), have been consistently shown to be increased in the cerebrospinal fluid (CSF) of ALS patients (7, 8).

The pivotal role of TDP-43 and other genes known to cause ALS (such as *FUS*, *hnRNPA2B1*, *hnRNPA1*, *TAF15* or *TIA1*) in the metabolism of RNA has implicated RNA dyshomeostasis as another crucial event in the pathophysiology of the disease (9). A comprehensive signature of transcriptional changes in the most affected ALS brain region is needed to further our understanding of this process. In this sense, an unbiased resource to obtain a comprehensive signature of gene and isoform expression changes associated with the whole tissue response to disease conditions is total RNA sequencing (RNAseq) in bulk tissue. Furthermore, the advent of novel sequencing tools such as single-nucleus RNAseq (snRNAseq) has opened a new window to better explore the transcriptome at a cell level, and has unraveled a complex and wide variety of human cell types with unique expression profiles in the human brain (10, 11, 12). These transcriptomic signatures can be applied to perform cell-type deconvolution of bulk RNAseq data using novel deconvolution methods. These methods take into account cross-subject and cross-cell variability of gene expression profiles, without relying on pre-selected markers, thereby providing more realistic cell-type proportion estimations and yielding crucial information about the cellular heterogeneity that is associated with a pathological condition (13).

The brain motor cortex is affected at the most early stages of the disease and is one of the most vulnerable regions in ALS. That being said, studies of this critical region both at the transcriptional and immunohistochemical level are lacking. In the present work, we aim to characterize the motor cortex of sporadic ALS cases through total RNAseq analyses. We also employ cell-type deconvolution using human snRNAseq data as reference and immunohistochemical analyses in order to resolve a signature of neuropathological changes associated with ALS in this vulnerable brain region.

## Material and methods

### Human samples

Human motor cortex (Brodmann Area 4) samples were provided by the Neurological Tissue Bank (NTB) of the Biobanc-Hospital Clínic-IDIBAPS. Clinical and neuropathological information of all ALS cases and healthy controls (HC) was carefully reviewed using data provided by the NTB. None of brain tissues presented any infarcts in the motor cortex. All ALS cases were clinically diagnosed with ALS, according to “El Escorial” criteria during life (14), had no family history of ALS or dementia, did not show any sign of cognitive impairment and presented pTDP43 inclusions in the motor cortex at postmortem examination. None of the samples presented the Samples carrying the *C9orf72* hexanucleotide repeat expansion or mutations in the *TBK1* gene, the most common adult-onset ALS gene mutations in Spain (15, 16).

### RNA extraction and sequencing

Using a mortar and liquid nitrogen, the tissue (60 mg) was grinded to powder and transferred to a solution of 600μl of TRIzol reagent (Thermo Fisher Scientific6). Standard recommended procedures were followed to extract RNA with TRIzol, the RNeasy mini kit (QIAGEN) and the Rnase free Dnase Set (QIAGEN). Qubit was used to measure RNA concentration whereas RNA integrity (RIN) was verified on an Agilent 2100 bioanalyzer (Agilent Technologies). Only samples with quantity threshold of 5 μg and RIN ≥ 6.5 were used for total RNAseq. The final study group included 11 ALS cases and 8 HC (Supplementary table 1). Paired-end sequencing libraries were prepared using the TruSeq Stranded Total RNA Library Preparation kit (Illumina) and sequenced on the Illumina HiSeq 2500 platform by the “Centro Nacional de Análisis Genómicos” (CNAG; Barcelona, Spain) with 101bp paired-end reads to achieve at least 100 million paired-end reads for each sample (ranging from 103.447.000 to 138.990.000 paired-end reads).

### Data availability

Raw sequencing data (FASTQ files) have been deposited at the European Genome-phenome Archive (EGA), which is hosted by the EBI and the CRG.

### Data processing

FASTQ files were aligned to the Grch38.p12 genome assembly using STAR v2.6.1a (17). In order to detect genetic variants, GATK Best Practices (18) for variant calling in RNAseq were followed. FeatureCounts (within the Subread v.1.6.2 package (19)) was used to assign fragments to each gene feature included in the Grch38v94 gene transfer format file. For differential isoform usage, STAR v.2.6.1a was used in “quantMode TranscriptomeSAM” mode and the output processed by Salmon (20) to obtain isoform expression values. Both differential gene expression and differential isoform usage analyses were performed using DESeq2 v1.24 (21). Gene ontology and KEGG pathway enrichment analyses were applied using default parameters (minimum overlap=3, p-value cutoff=0.01 and minimum enrichment=1.5) in Metascape 3.0 (22) (http://metascape.org). Synaptic enrichment analyses were also run using the web server SynGO (https://www.syngoportal.org/), which provides an expert-curated resource for synapse function and gene enrichment analysis (23).

### Validation of transcriptome changes by qPCR

Quantitative real-time PCR (qPCR) was performed on the same brain-derived RNA samples used for total RNAseq. A total of 500ng of RNA was used to generate cDNA with the RevertAid First Strand cDNA Synthesis Kit (ThermoFisher), as per manufacturer’s instructions. All qPCRs were conducted using Fast SYBR Green Master Mix (ThermoFisher) and run on an ABI Prism 7900HT Fast Real-Time PCR System (Applied Biosystems). All primer pairs used are listed in supplementary table 5. Relative quantification was determined using the ΔΔCt method and normalized to the endogenous control *RPLP13*.

### Weighted Gene Co-expression Network Analyses (WGCNA)

Raw gene counts were filtered and only genes with a minimum of 10 counts across all individuals were kept for further analyses. DESeq2 was then used to normalize the input data to construct co-expression networks using the WGCNA (24). The signed weighted correlation network, which takes into account both negative and positive correlations, was constructed using the manual function in the WGCNA package, using the “biweight midcorrelation”, selecting a power of 7 (the lowest possible power term where topology fits a scale free network) and run in a single block analysis. Module definitions were calculated by the hybrid treecutting option with “deepsplit parameter” = 2. Minimal module size was set at 30 and modules with a cutHeight ≤ 0.25 were merged. Modules were summarized by module eigengenes (ME) and correlations to traits were performed with spearman correlation for continuous traits and Pearson correlation for binary traits (bicor and corPvalueStudent, included in the WGCNA package). Module eigengenes (first principal component as a summary of this module) were calculated for each individual and module to perform correlation analyses.

### Cell-type deconvolution

The recently developed Multi-Subject Single Cell deconvolution method (MuSiC) (13) was used to perform cell-type deconvolution of our bulk RNAseq data. As a reference, we used the most comprehensive human snRNAseq dataset available. This dataset includes the RNAseq from 70.634 cells in the frontal cortex (Brodmann Area 10) of 24 Alzheimer’s disease cases and 24 HC from Religious Orders Study (ROS) and Memory and Aging Project (MAP) (ROSMAP) (12). This data file comprises 70.634 cells derived from the frontal cortex (Brodmann Area 10) grouped into 8 major cellular populations (excitatory neurons, inhibitory neurons, microglia, astrocytes, oligodendrocytes, oligondendrocyte precursor cells, endothelial cells and pericytes). In order to check consistency and validate our results, major cell-type proportions were identified using two other snRNAseq repositories as a reference. We downloaded the medial temporal gyrus snRNAseq data derived from the Allen Brain Atlas, which includes 15.928 cells from 8 human tissue donors. We selected this dataset as it contains double the number nuclei than the primary visual cortex and the anterior cingulate cortex included in the Allen Brain Atlas (10). Finally, the third dataset included 10.319 cells derived from the frontal cortex of 4 individuals (11). Cell-type deconvolution of cellular subpopulations was performed using the ROSMAP data as reference.

### Immunohistochemistry

Formalin-fixed pParaffin-embedded (FFPE) motor cortex tissue sections of 5μm were analyzed by immunohistochemistry (IHC) (n=11 ALS patients, n=7 controls). One control sample was excluded from the analysis due to lack of availability of motor cortex sections. The FFPE sections were dewaxed, by placing them in xylene and decreasing concentrations of ethanol, and hydrated 30 minutes in distilled water. Tissue sections were treated with a solution containing 10% methanol, 3% H_2_O_2_ and PBS for 10 minutes, to inhibit endogenous peroxidase. These sections were pre-treated and boiled in citrate 1x solution to mediate antigen retrieval. Once cooled, sections were blocked with 5% bovine serum albumin for 1 hour at room temperature. Sections were incubated overnight with the corresponding primary antibody at 4°C and then incubated with the anti-rabbit horse radish peroxidase secondary antibody (1:200) for 1 hour at room temperature. Primary antibodies were Iba1 rabbit policlonal (1:500; Wako Chemicals) and ps409/410-TDP rabbit polyclonal (1:2000; Cosmo Bio). Following peroxidase development with 3,3′-diaminobenzidine tetrahydrochloride (Dako liquid DAB + substrate Chromogen System, DakoDenmarck), sections were stained with hematoxylin (EnVision™ FLEX haematoxylin, Dako Denmark) and dehydrated using increasing concentrations of ethanol and then mounted with DPX mounting medium (PanReacApplichem, ITW Reagents). For immunofluorescence, 3 ALS and 3 control motor cortex sections were incubated overnight at 4°C in primary antibody. Primary antibodies were Iba1 rabbit policlonal (1:500; Wako Chemicals) and Human HLA-DP, DQ, DR antigen, mouse clone CR3/43 (1:20, Agilent, M077501–2). Sections were incubated for 1 hour at room temperature with secondary antibodies (1:1000). Secondary antibodies were conjugated with Alexa-Fluor 488 and Alexa-Fluor 555 (Invitrogen), and cell nuclei visualized with Hoescht 33258 (Life Technologies). Sections were mounted with Shandon™ Immu-Mount™ (Thermo Scientific). Images were acquired using a confocal microscope (Leica TCS SP5 Confocal Laser Scanning Microscope) with a 63× objective.

### Quantification analyses

Full-section IHC images were obtained with Pannoramic MIDI II (3DHistech). Six grey matter areas of motor cortex from each case were delimited blinded to neuropathological diagnoses. An in-house computer-based algorithm was developed to quantify densities for Iba1 staining. The algorithm-based approaches were performed with the MATLAB R2017b software (The MathWorks, Inc., Natick, MA, USA). For pTDP43 quantification, immunoreactive positive objects were quantified manually by two different researchers blinded to neuropathological diagnoses.

### Statistical Analyses

Shapiro-Wilk test was used to test for deviation from a standard distribution. Correlation analyses were performed using Pearson’s or Spearman’s test (using the *cor.test* function in R), as well as Student-t test or Student-test with Welch correction (using *t.test* function in R) to compare means, depending on data distribution. A p<0.05 was considered significant.

## Results

Postmortem motor cortex samples from 11 ALS cases (6 females) and 8 subjects without neurological disease (4 females) were analyzed by RNAseq. Postmortem interval and RIN did not differ between groups (p=0.68 and p=0.089, respectively). The mean age at death was lower for ALS cases compared to HC (63.4 years; range: 54-82 years in ALS and 73.9 years; range: 58-83 years in HC, *p*=0.04). Patients had an average disease onset of 60.9 years (range: 46-81) and a disease duration of 29,9 months (range: 1-72). Limb onset was presented in nine out of the eleven cases (81.8%) (Supplementary table 1). Using RNAseq data, the presence of other ALS-disease causing mutations were ruled out in all samples, confirming the presumably sporadic nature of our ALS group of cases.

Differential gene expression between ALS patients and HC disclosed a total of 108 up-regulated and 16 down-regulated genes (adj. *p*<0.05) (figure 1A and supplementary table 2). Altered expression was validated by qPCR for nine out of the ten genes selected. Only the *CHI3L2* gene presented a non-significant trend towards an overexpression in ALS patients (*p*=0.085) (figure1B). Correlation analyses between counts derived from total RNAseq data and qPCR expression values indicated a high and significant concordance between both expression measures in the 10 genes that were evaluated (supplementary table 3). Gene ontology (GO) and KEGG pathway analyses revealed an enriched involvement of the inflammatory response in ALS (figure 1C). The assessment of changes at the isoform expression level resulted in 167 upregulated and 40 downregulated isoforms, of which 181 were from unique genes (supplementary table 4). Enrichment analyses showed the postsynaptic density as the main altered GO term and the immune response as dominant pathways (supplementary figure 1). To further characterize and confirm the synaptic involvement, we used the SynGO database, which provides an expert-curated resource for synapse function and gene enrichment analysis (23). This analysis confirmed the enrichment of postsynaptic components in our list of differentially expressed isoforms (supplementary table 5).

**Figure 1.**
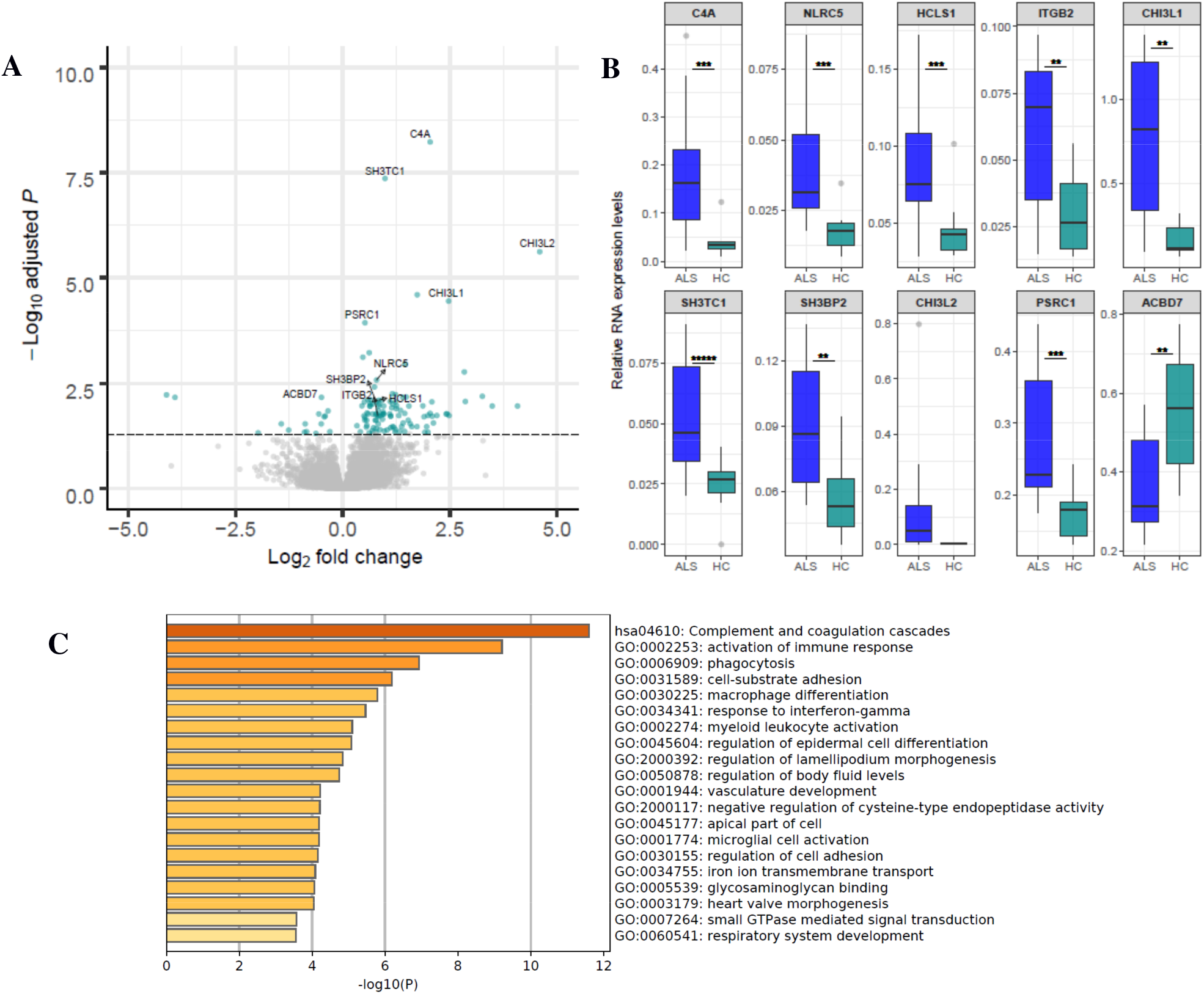
Differential gene expression changes in the sporadic ALS motor cortex. Volcano plot displaying the significant differentially expressed genes between the ALS and the HC motor cortex. The vertical axis (y-axis) corresponds to the −log_10_ adjusted p-value and the horizontal axis (x-axis) represents the log_2_ fold change value. Blue circles correspond to the significantly differential genes expressed in this tudy (adjusted p-value<0.05), whereas grey circles display the non-significant genes. Gene names are shown for the ten genes selected for qPCR validation (A). Bar chart showing the relative RNA expression values in ALS and HC obtained through qPCR for the ten genes selected for validation. *p<0.05; **p<0.01; ***p<0.001 and ***** p<0.00001 (B). Gene ontology and KEGG pathway enrichment analyses obtained from the list of genes showing a significant differential expression (C).

In order to find clusters of highly correlated genes, we next performed weighted gene co-expression network analyses (WGCNA), which disclosed a total of 8 modules. Among them, three modules significantly correlated with disease status (MEblack, *p*=0.003; R=0.64; MEyellow, *p*=0.025; R=0.57 and MEpink, *p*=0.011; R=−0.51). Moreover, MEblack and MEyellow were enriched for inflammatory responses, whereas MEpink had an over representation of genes involved in synaptic and neuronal functions (table 1). The SynGO database confirmed the postsynaptic signature of this enrichment in MEpink (Supplementary table 6). Interestingly, the two inflammatory gene co-expression modules (MEblack and MEyellow) showed a strong negative correlation with the postsynaptic module (MEpink) (*p*=1.3×10^−4^; R=−0.77 and *p*=9×10^−4^; R=−0.69, respectively) (supplementary figure 2), suggesting that both inflammation and synaptic alterations are interconnected in ALS pathological processes.

**Table 1.** Gene expression modules obtained by weighted gene co-expression analyses significantly associated with ALS. The three most significant gene ontology terms and their enrichment log_10_ p-value are shown for each module.

|  | P value | R | Number of genes | Most significant GO term | Log10P enrichment |
| --- | --- | --- | --- | --- | --- |
| **MEblack** | 0.003 | 0.637 | 439 | myeloid leukocyte activation | -33.41 |
|  |  |  |  | activation of immune response | -28.84 |
|  |  |  |  | cytokine production | -25.89 |
| **MEpink** | 0.011 | -0.512 | 851 | postsynaptic membrane | -13.54 |
|  |  |  |  | potassium channel activity | -8.59 |
|  |  |  |  | neuron projection morphogenesis | -6.06 |
| **MEyellow** | 0.025 | 0.572 | 1145 | cytokine-mediated signaling pathway | -19.94 |
|  |  |  |  | adherens junction | -18.18 |
|  |  |  |  | blood vessel development | -17.16 |

To characterize the variability of cell types and compare cellular proportions between groups, we applied the recently developed MuSiC algorithm. As reference, we used data from the first and most comprehensive human snRNAseq study performed in brain tissue of subjects with a neurodegenerative disease which included 2 subjects with Alzheimer’s disease and 24 HC (12). Cell-type deconvolution revealed an overrepresentation of microglial cells in the ALS motor cortex as compared to HC (*p*=0.025) and a trend towards a reduction in excitatory neurons (*p*=0.101) (figure 2). Microglial upregulation was confirmed using two independent reference datasets: the Allen Brain Atlas and Lake and collaborators (10, 11). In both cases, the same trend was found towards the down-regulation of excitatory neurons (supplementary figure 3 and 4). Importantly, the estimation of major cellular populations significantly correlated between the three datasets (except for inhibitory cells, as these correlations only emerged when comparing the datasets from Lake and collaborators (11) and Allen Brain atlas (10)), thus strengthening our deconvolution-based results. To further gain insight into which microglial subpopulation might be driving this effect, we estimated the proportion of each subpopulation identified in the ROSMAP study. Our results showed that a specific microglial subpopulation (Mic1, as named in the study performed by Mathys and collaboratorsand recognized as the human DAM (12)) is disproportionately presented in the ALS motor cortex (*p*=6.6×10^−3^) (figure 3A). Among the 77 marker genes which characterize the Mic1 subcluster as DAM (Mic1 unique genes or those overlapping with DAM) (11), 22 of them (28.6%) are genes differentially expressed in our bulk RNAseq study (unadjusted *p*<0.05; (Supplementary table 7). Furthermore, the proportion of Mic1 cells showed a strong positive correlation with the expression of *TREM2* (p=6.8×10^−4;^; R=0.71), required to trigger the DAM profile, and the two inflammatory gene co-expression modules associated with ALS (Meblack; *p*=2.7×10^−8^, R=0.92; and Meyellow; *p*=1.4×10^−3^, R=0.68). A negative was also found between Mic1 and the synaptic module (MEpink; *p*=2.4×10^−4^, R=−0.75) (figure 3B), but did not correlate with the other gene co-expression modules identified in this study. Overall, our results from differential gene and isoform expression comparisons, gene co-expression modules and cell-type deconvolution, strongly indicate that microglia-related inflammatory changes, mainly driven by the Mic1 subpopulation, are central in ALS pathophysiology and may be closely related to the synaptic changes that are present in the motor cortex.

**Figure 2.**
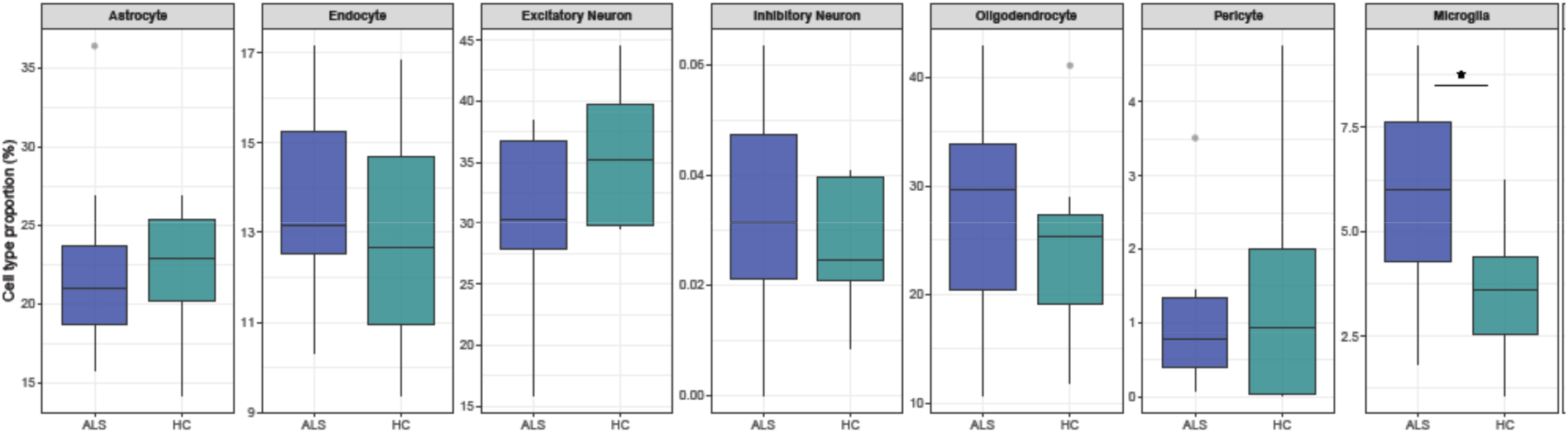
Cell type composition in the ALS and HC motor cortex. Box-plot showing the estimated major cell type proportions for the ALS and HC by MuSiC obtained using the human single-nucleus RNAseq (snRNAseq) data from 24 Alzheimer’s disease and 24 healthy controls (ROSMAP) human frontal cortex (Brodmann Area 10). *<0.05

**Figure 3.**
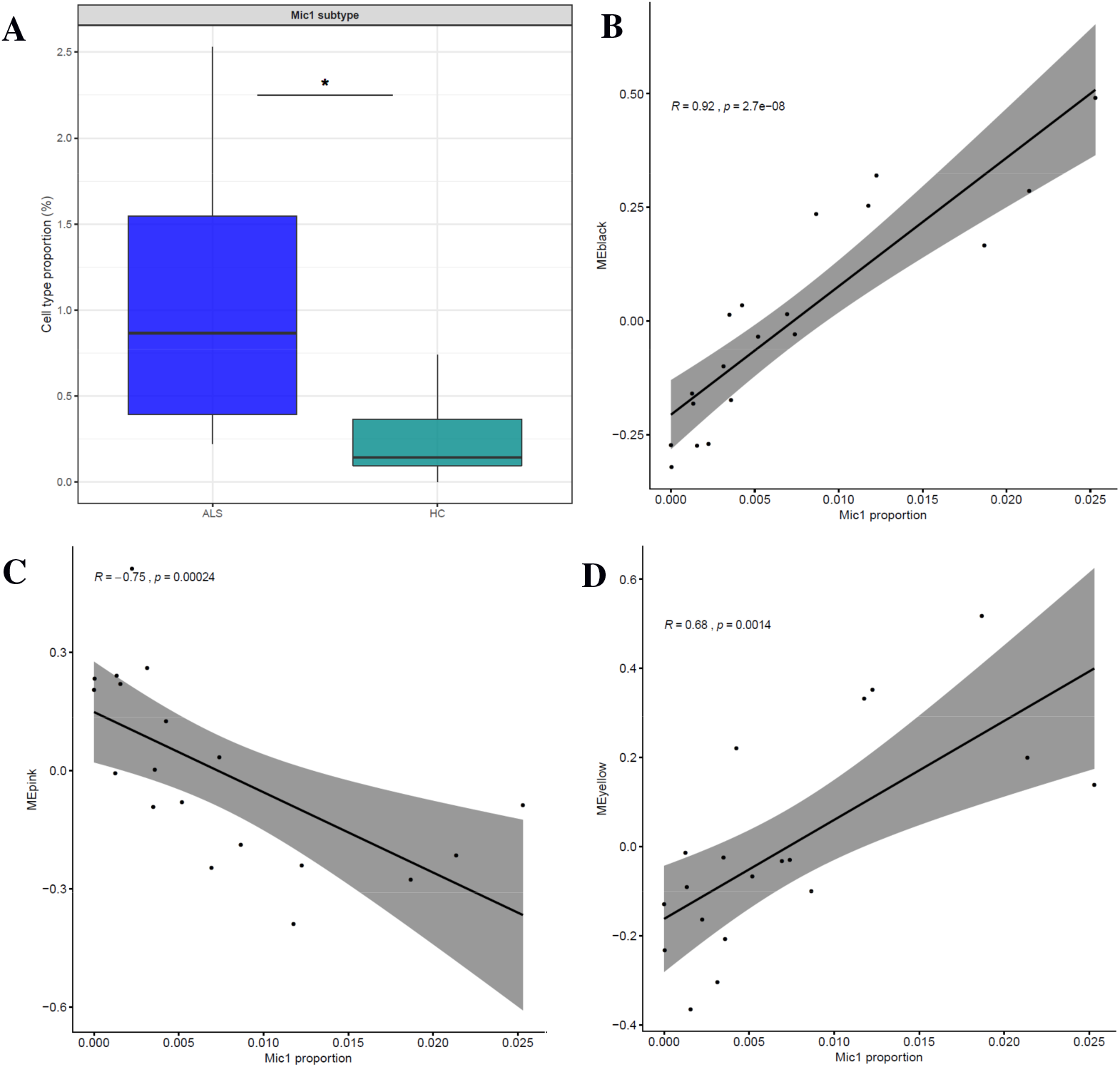
Estimation of the Mic1 microglial subpopulation proportion identified using the ROSMAP human snRNAseq dataset (A). Correlation analyses between Mic1 proportion and the individual eigengenes obtained from the MEblack (B), MEpink (C) and MEyellow (D) gene expression modules. \*\**p*<0,01

To further investigate our results in postmortem brain tissue, we performed IHC analyses using Iba1 to examine the broad population of microglial cells, and MHC class II as a marker of reactive microglia that is expressed in the Mic1 cell subpopulation. Positive immunostaining was increased in the ALS motor cortex for Iba1, however, differences between patients and controls did not reach statistical significance (*p*=0.106). Notwithstanding, co-immunofluorescence of Iba1 and MHC class II markers revealed an increase in the number of microglial cells expressing MHCII markers in ALS as compared to HC motor cortex (figure 4), as previously described (12), strengthening our results and suggesting that Mic1 is driving neuroinflammation in the ALS motor cortex.

**Figure 4.**
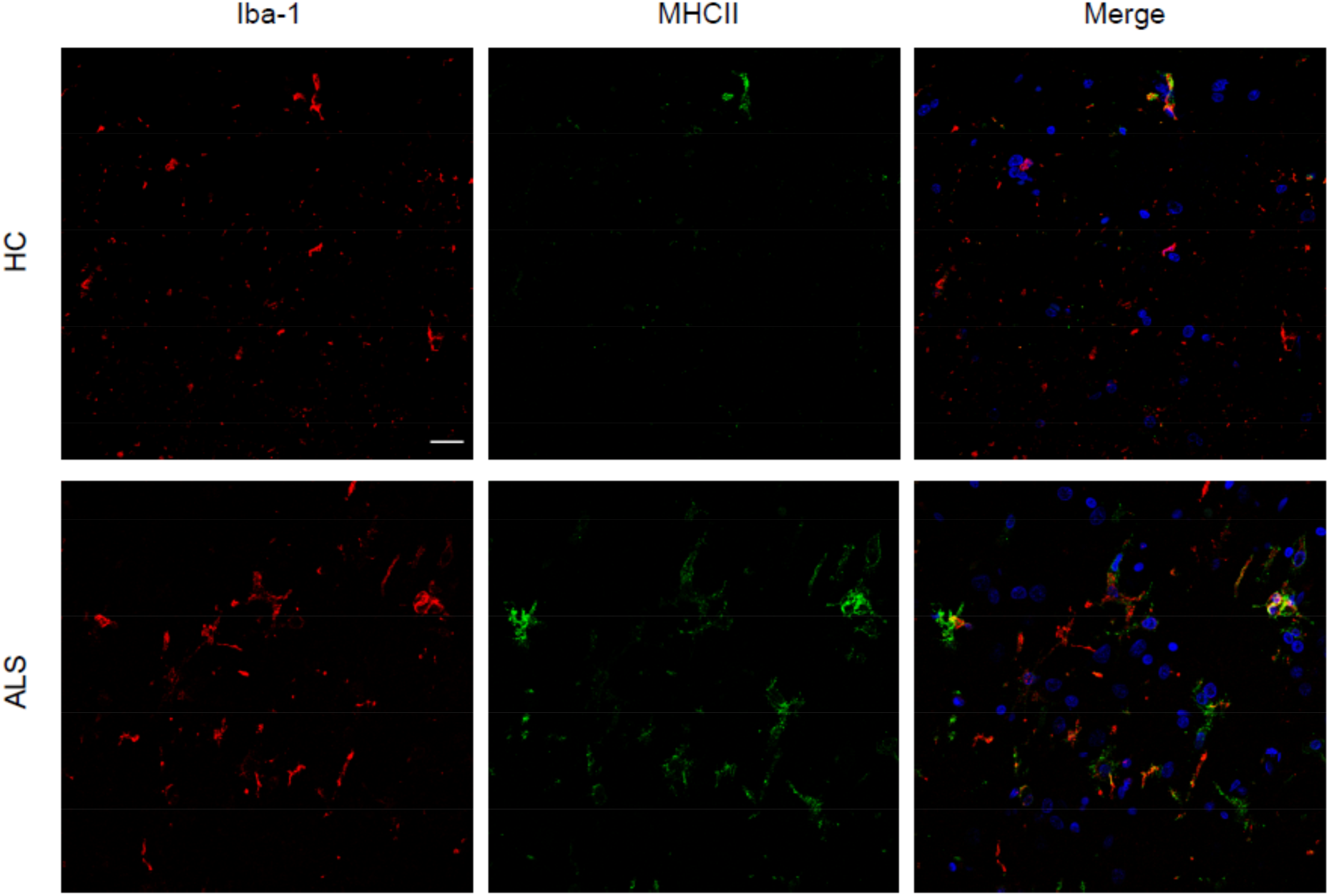
Co-immunofluorescence with anti-IBA1 (red) and anti-MHC class II (green) antibodies in the ALS and HC motor cortex.

We finally assessed the frequency of pTDP43 immunoreactive structures in the motor cortex. As expected, the density of pTDP43 inclusions was increased in ALS as compared to HC (*p*=0.017) (figure 5A). Interestingly, a high degree of variability on the number pTDP43 aggregates was noted in ALS, and was not explained by age at death, age at onset or disease duration. The burden of pTDP43 inclusions correlated inversely with the proportion of microglial cells in our group of ALS patients (*p*=0.016; R=−0.7) (figure 5B). Notably, the same trend was found with the microglial subpopulation Mic1 (*p*=0.071; R=−0.56). No correlations were detected with any of the other major cell types nor microglial subtypes.

**Figure 5.**
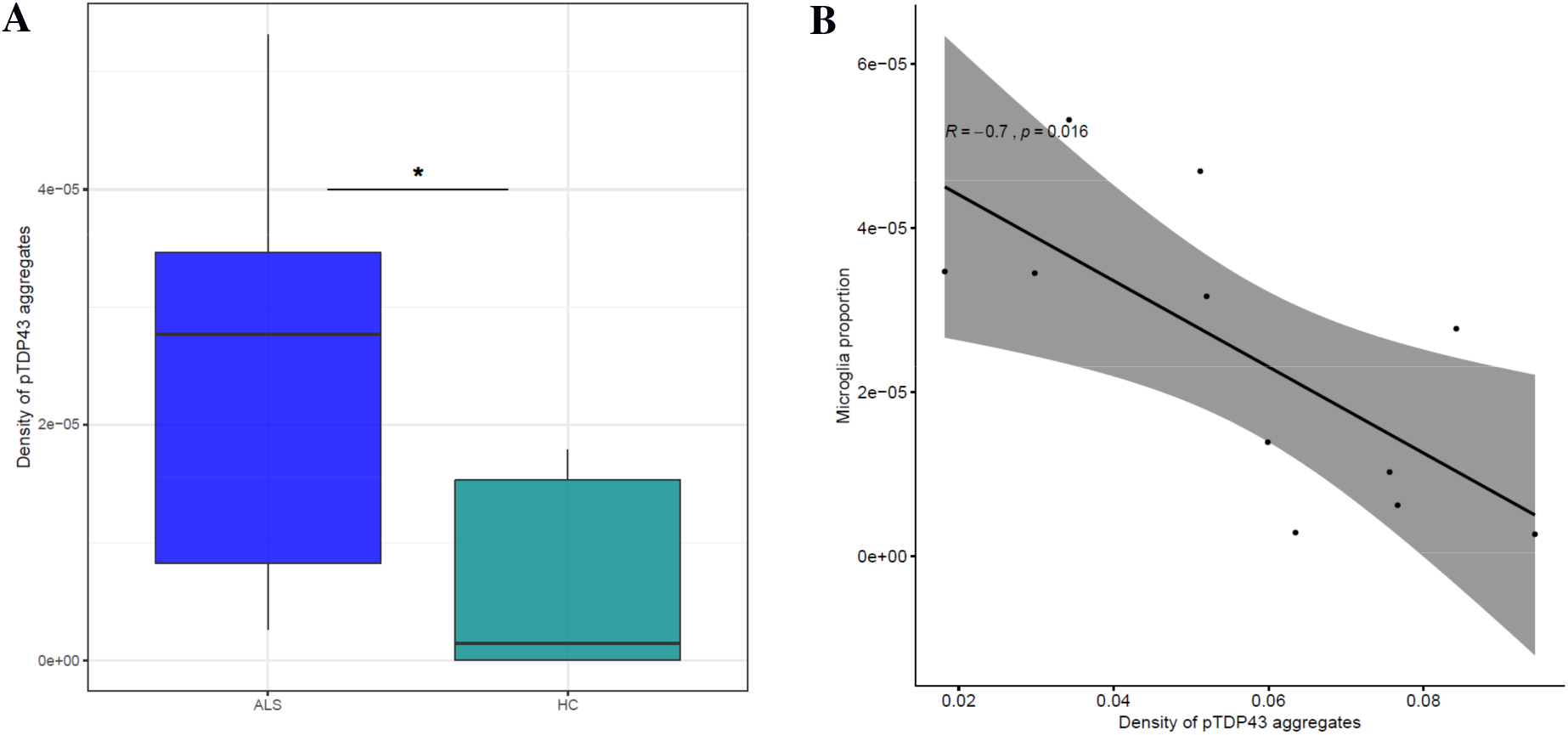
Analysis of the burden of pTDP43 aggregates in the ALS and HC motor cortex. Density of pTDP43 aggregates in each postmortem individual included in this study (A). The burden of pTDP43 aggregates correlates with the estimates of microglial proportion in the ALS group of cases (B).

## Discussion

In order to better understand the biological bases of sporadic ALS, we decided to investigate perturbations in RNA expression, one of the most prominent pathogenic mechanisms in ALS (9), in the motor cortex of ALS patients not harboring an ALS-causing mutation, without family history of ALS or dementia and with no signs of cognitive impairment in the retrospectively reviewed clinical files. As the motor cortex is one of the major vulnerable and early affected regions in the disease, it represents a target region to disentangle key pathological processes in this neurodegenerative disorder. To date, few studies have performed a whole transcriptomic assessment of the CNS in ALS (25, 26, 27). However, this is the first deep transcriptome characterization of the ALS motor cortex with a multimodal approach that encompasses total RNAseq, cell-type deconvolution using human snRNAseq and immunohistochemical assessments. We report a list of 124 genes differentially expressed in the ALS motor cortex which reflect the RNA expression changes in this brain region. Among them, chitinase-related genes *CHI3L1* and *CHI3L2* represent two of the most prominent shifts in gene expression, whereas *CHIT1* shows a nominal significance. Interestingly, these chitinases are neuroinflammatory biomarkers which have been previously shown to be consistently increased in CSF of ALS patients compared to neurologically healthy subjects (7, 8). These results indicate that, among the unbiased signature of RNA alterations provided herein, some of them might be promising biomarkers that require further investigation.

An exacerbated innate immune response, with microglial activation as a predominant signature, has been recently shown in the ALS motor cortex (28). Furthermore, the only study which has investigated the whole transcriptome of the ALS motor cortex suggests that, among ALS cases, a subgroup of them is characterized by a higher inflammatory response (27). Our results also indicate that microglial-related neuroinflammatory changes in the human motor cortex are critical in the pathophysiology of this neurodegenerative disorder. First, differential gene and isoform expression data points towards an inflammatory response as a major event that is significantly intensified in the ALS motor cortex. Second, the assembly of weighted gene co-expression networks resulted in two significant modules associated with ALS (MEblack and Meyellow), both highly enriched with genes associated with inflammatory functions. Third, cell-type deconvolution of the bulk RNAseq data demonstrated an increased proportion of microglial cells as compared to the motor cortex of healthy cases. In this context, studies in mice have recently suggested DAM as the microglial subpopulation with a more prominent role in ALS (4). For the first time in the context of human motor cortex, our results show that Mic1, the human microglial subpopulation which harbors the majority of markers found in the previously described DAM transcriptomic signature (12), is the main microglial subpopulation which drives microgliosis in ALS and that is heavily correlated with the expression of *TREM2*, which is required to enhance the late and higher pro-inflammatory stage of DAM (4).

Our immunohistochemical analyses did not show a relevant increased density of Iba1 (a widely used marker of microglial processes) in ALS-related brain tissue. Previous studies have frequently provided contradictory results when using this marker to assess microgliosis (29). This somewhat unexpected result might be explained by the fact that, whereas cell-type deconvolution is performed using a complete catalogue of gene expression profiles obtained from brain-derived snRNAseq, immunohistochemistry only uses a single marker and does not reflect the complexity and heterogeneity of this cell type, making it an underpowered method to detect differences in the proportion of cell subpopulations. That being said, we did validate the increased proportion of Mic1 cells within the ALS motor cortex through co-immunofluorescence of two markers (Iba1 and MHC class II) previously shown to characterize this microglial subpopulation (12). Thiss finding further establishes Mic1 (the human “disease associated microglia”) as the microglial subpopulation that drives microgliosis in the ALS motor cortex.

We observed a high degree of variability in the the density of pTDP43 inclusions across patients, which could not be explained by any of the demographic or clinical features available in this study. Notably, we found a striking inverse correlation between the proportion of microglial cells and the amount of pTDP43 aggregates. These results are in line with recent *in vivo* and *in vitro* studies, suggesting that although microgliosis arises as a phagocytic response to pTDP43 aggregates, at some point these cells lose their ability to clear these neuropathological insults and are downregulated (30, 31). Whether biofluid levels of microglial markers, such as TREM2, could be used as a proxy of pTDP-43 density in the motor cortex is an avenue worth pursuing and would be a valuable addition to the biomarker arsenal for use in designing clinical trials and assessing therapeutic efficacy.

Synaptic dysfunction is an early pathogenic event in ALS (32). Our data point towards an over-representation of postsynaptic markers and a trend towards a decrease of excitatory neurons in ALS patients. Together, these results are consistent with upper motor neuron degeneration occurring in this brain region. Recent studies in other neurodegenerative diseases have suggested that microglia is a key and early mediator of synapse loss through phagocytosis induced by the complement cascade (33, 34). In fact, our gene expression data show an increased expression of some complement cascade related genes, reinforcing this hypothesis. Furthermore, our results indicate that the two microglial-related neuroinflammatory modules of gene expression (MEblack and MEyellow) as well as the proportion of Mic1 cells inversely correlate with the presence of synaptic markers, suggesting that activated microglia and synapse loss in the motor cortex are highly related processes in ALS pathogenesis.

Overall, our study strongly reinforces the key role of microglia and neuroinflammation in ALS, being the DAM subpopulation a key element driving this effect. The identification of specific microglial populations with well-defined transcriptional signatures will contribute to disentangle new mechanisms and novel therapeutic targets to fight against this devastating disorder.

## Supporting information

Supplementary table 1. Demographic and clinical data of ALS and HC included in this study. RIN: RNA integrity number. PMI: Postmortem interval.

Supplementary table 2. List of the genes showing a significant differential expression in the ALS motor cortex as compared to HC provided by DESeq2.

Supplementary table 3. Correlation analyses between gene expression counts obtained through RNAseq and RNA relative expression values derived from qPC

Supplementary table 4. List of the isoforms showing a significant diferential usage in the ALS motor cortex as compared to HC provided by DESeq2.

Supplementary table 5. Enrichment of synaptic markers within the list of differently expressed isoforms using SynGO ontology terms.

Supplementary table 6. Enrichment of synaptic markers within the MEpink gene expression module using SynGO ontology terms.

Supplementary table 7. List of genes differentially expressed (unadjusted p-value<0.05) in the ALS motor cortex provided by DESeq2 which overlap with

Supplementary figure 1. Gene ontology and KEGG pathway enrichment analyses obtained from the list of isoforms showing a significant differential usage

Supplementary figure 2. Correlation analyses between the Meblack and Meyellow (enriched with inflammatory markers) and the MEpink (enriched with synap

Supplementary figure 3. Estimation of cell-type proportions by MuSiC using the Lake et. al, human single-nucleus RNAseq dataset; *p<0.05.

Supplementary figure 4. Estimation of cell-type proportions by MuSiC using the Allen Brain Atlas human single-nucleus RNAseq dataset; *p<0.05.

## Aknowledgements

We are indebted to the IDIBAPS Biobank for sample and data procurement.

