## Supplementary figures and images for "Characterization of the motor cortex transcriptome supports microgial-related key events in amyotrophic lateral sclerosis"

### Supplementary figure 1. Gene ontology and KEGG pathway enrichment analyses obtained from the list of isoforms showing a significant differential usage

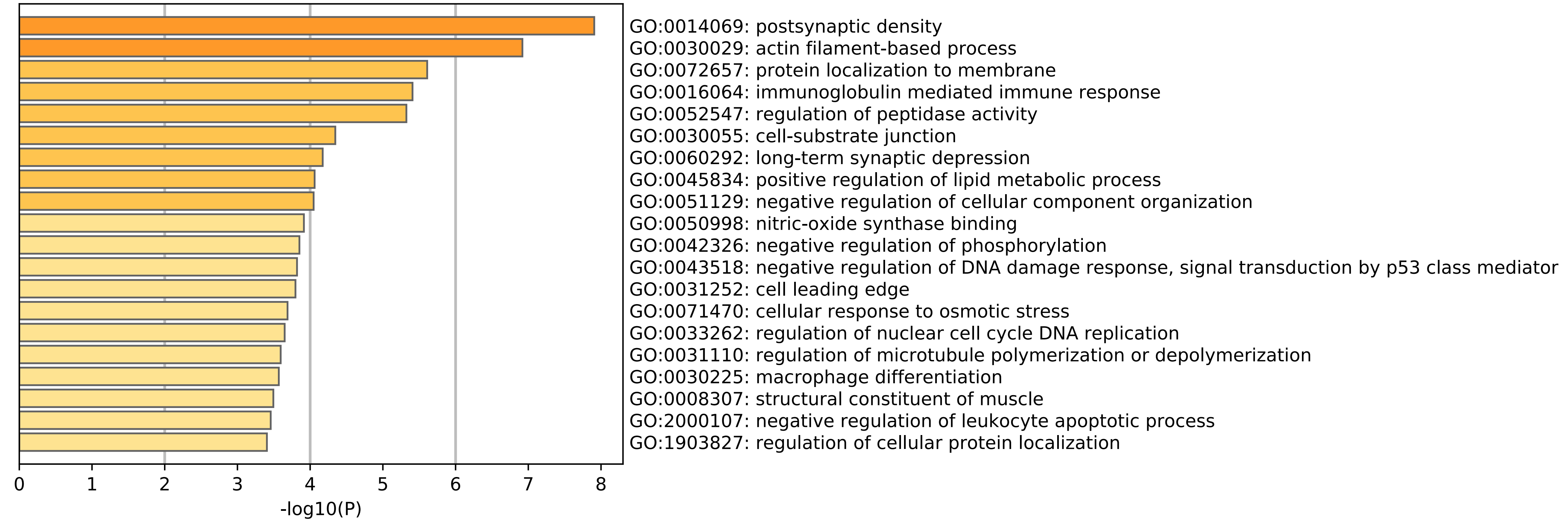

### Supplementary figure 2. Correlation analyses between the Meblack and Meyellow (enriched with inflammatory markers) and the MEpink (enriched with synap

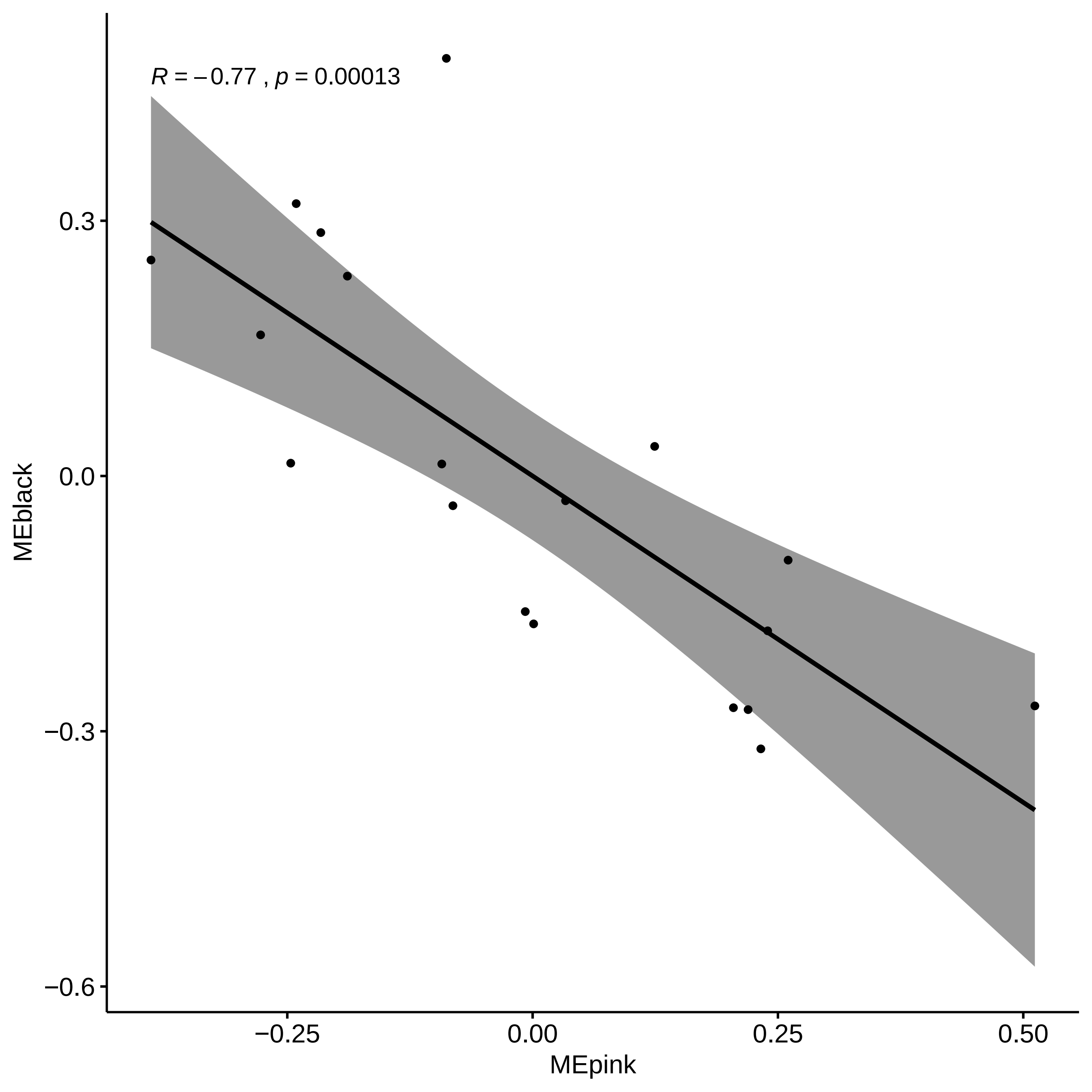

### Supplementary figure 3. Estimation of cell-type proportions by MuSiC using the Lake et. al, human single-nucleus RNAseq dataset; *p<0.05.

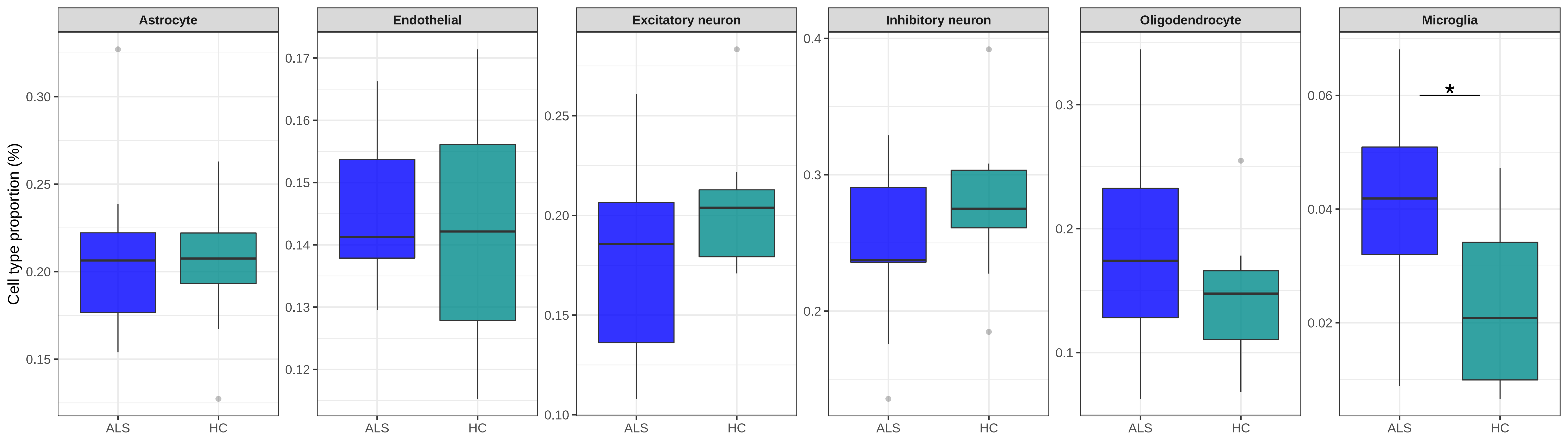

### Supplementary figure 4. Estimation of cell-type proportions by MuSiC using the Allen Brain Atlas human single-nucleus RNAseq dataset; *p<0.05.

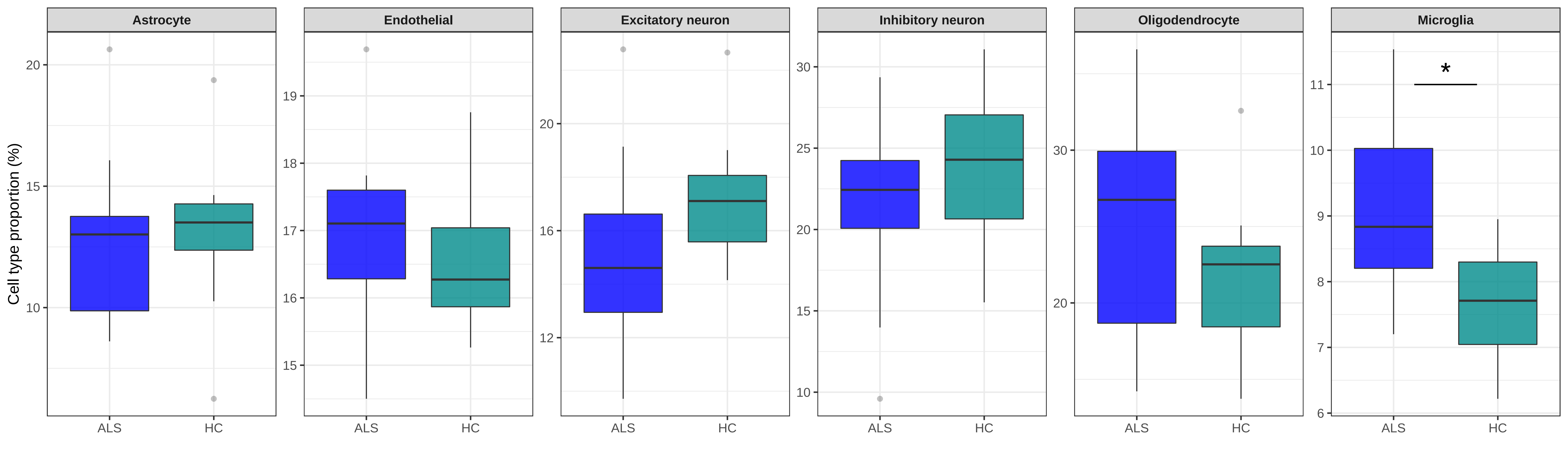
